# Physiological Changes Associated with Copper Sulfate-Induced Emesis in Felines

**DOI:** 10.1101/2022.10.20.512908

**Authors:** Charles P. Murphey, Jonathan A. Shulgach, Pooja R. Amin, Nerone K. Douglas, John P. Bielanin, Jacob T. Sampson, Charles C. Horn, Bill J. Yates

**Author notes:** **Correspondence:** Bill J Yates.

## Abstract

Nausea is a common disease symptom, yet there is no consensus regarding its physiological markers. In contrast, the process of vomiting is well documented as sequential muscular contractions of the diaphragm and abdominal muscles and esophageal shortening. Nausea, like other self-reported perceptions, is difficult to distinguish in preclinical models, but based on human experience emesis is usually preceded by nausea. Here we focused on measuring gastrointestinal and cardiorespiratory changes prior to emesis to provide additional insights into markers for nausea. Felines were instrumented to chronically record heart rate, respiration, and electromyographic (EMG) activity from the stomach and duodenum before and after intragastric delivery of saline or copper sulfate (CuSO_4_, from 83 to 322 mg). CuSO_4_ is a prototypical emetic test agent that triggers vomiting primarily by action on GI vagal afferent fibers when administered intragastrically. CuSO_4_ infusion elicited a significant increase in heart rate, decrease in respiratory rate, and a disruption of gastric and intestinal EMG activity several minutes prior to emesis. The change in EMG activity was most consistent in the duodenum. Administration of saline did not induce these effects. Increasing the dose of CuSO_4_ did not alter the physiologic changes induced by the treatment. It is postulated that the intestinal EMG activity was related to a retrograde movement of chyme from the intestine to the stomach. These findings suggest that monitoring of intestinal EMG activity, perhaps in combination with heart rate, may provide the best indicator of the onset of nausea following treatments and in disease conditions, including GI disease, associated with emesis.

## 1 Introduction

Nausea is ubiquitous in many disease states, including gastrointestinal (GI) disease and vestibular disorders, and it is a common side effect of many clinical treatments, such as cancer chemotherapy, inhalational anesthesia, and opioid pain medications (Golding and Gresty, 2015; Piechotta et al., 2021; Weibel et al., 2021). Nausea is also difficult to treat, especially in chronic conditions where patients are often provided anti-emetics, which are less effective for the control of nausea (Koch et al., 2016; Lacy et al., 2018). In contrast to other conditions, such as pain and anxiety, the physiological indicators of nausea are not well established. Based on human experience, emesis, which is produced through a well-characterized sequence of muscle contractions (Grelot and Miller, 1994), is usually preceded by nausea.

Changes in cardiac and GI myoelectric responses are commonly reported correlates of nausea in humans, specifically related to emetogenic platinum-based chemotherapy (using drugs such as cisplatin), GI diseases, and motion sickness (Morrow et al., 2000; Himi et al., 2004; Carson et al., 2022). Increased heart rate, decreased heart rate variability, and changes in gastric myoelectric activity have been reported to be associated with nausea (Morrow et al., 2000; Himi et al., 2004; Kim et al., 2011; LaCount et al., 2011). Several preclinical studies using the most common emetic test species (dog, cat, and ferret) have tracked the temporal associations between emesis and these responses. In dogs, cisplatin chemotherapy and intragastric copper sulphate (CuSO_4_) disrupt intestinal and gastric myoelectric activity (Lee et al., 1985; Chey et al., 1988; Ando et al., 2014). In cats, centrally acting emetic agents, particularly morphine and apomorphine, disrupt intestinal myoelectric activity associated with emesis (Stewart et al., 1977). In addition, studies using ferrets showed simultaneous changes in cardiac and GI myoelectric measures following emetic treatments; for example, cisplatin treatment produced gastric myoelectric disruption, an increase in heart rate, and a decrease in heart rate variability (Lu et al., 2017). Limitations of these prior studies include a lack of simultaneous measures in each experiment (e.g., intestinal, gastric, and cardiorespiratory recordings). In addition, most failed to use an acute emetic stimulus, such as intragastric CuSO_4_, to permit temporal associations between physiological measures and the onset of emesis.

The focus of the current study was to examine changes in cardiorespiratory, gastric and intestinal myoelectric signals preceding emesis, and, therefore, to ascertain potential markers for nausea. We used the cat model in the current work because of its well-documented physiologic responses during emesis (Makale and King, 1992; Hickman et al., 2008), ease of surgical electrode placement, and adaptability to behavioral training. Importantly, we trained animals to remain sedentary while loosely restrained to limit the occurrence of electrical artifacts. We also used intragastric CuSO_4_ as the emetic test stimulus because copper and its derivatives are known to elicit nausea in humans (Pizarro et al., 1999; Araya et al., 2001). CuSO_4_ rapidly triggers emesis in cats and other preclinical models primarily by activating the GI vagus nerve-to-brain pathway (Wang and Borison, 1951; Brizzee and Marshall, 1960; Makale and King, 1992; Horn et al., 2014). The acute action of this agent allowed us to monitor a variety of physiologic parameters prior to emesis during a controlled recording session. Before and following CuSO_4_ treatment we recorded heart rate, respiration rate, and gastric and duodenal myoelectric responses using methodology developed through our work in ferrets (Nanivadekar et al., 2019; Shulgach et al., 2021). The dose of CuSO_4_ was additionally varied between animals so we could determine whether increasing the dose altered physiologic responses prior to emesis.

## 2 Methods and Materials

### Animals and Husbandry

All experimental procedures on animals were approved by the University of Pittsburgh’s Institutional Animal Care and Use Committee, in accordance with the National Research Council’s *Guide for the Care and Use of Laboratory Animals* (National Research Council, 2011). Experiments were performed on 10 antibody profile defined and specific pathogen free domestic shorthair cats (6 male, 4 female) obtained from Marshall BioResources (North Rose, New York, USA). Animals were provided commercial cat food and water *ad libitum* and were housed under 12h light/dark cycles. Characteristics about the animals and other experimental parameters are provided in *Table 1*.

**Table 1.** Characteristics of animals used in the experiments, as well as experimental parameters.

| **Animal ID** | **Sex** | **# Acclimation Days Prior to Surgery** | **# Days Following Surgery and Prior to Final Experiment** | **Weight at Final Experiment (Kg)** | **Total Number of Recording Sessions (Control + Final Experiment)** | **CuSO<sub>4</sub> Dosage (mg)** | **Method of CuSO<sub>4</sub> Administration** |
| --- | --- | --- | --- | --- | --- | --- | --- |
| C1-21 | F | 65 | 48 | 3.78 | 29 | 163 | Intragastric Catheter |
| C2-21 | F | 92 | 30 | 3.44 | 11 | 332 | Oral |
| C3-21 | F | 51 | 57 | 2.75 | 32 | 332 | Oral |
| C11-21 | F | 36 | 43 | 3.70 | 23 | 332 | Oral |
| C13-20 | M | 154 | 50 | 4.70 | 23 | 83 | Intragastric Catheter |
| C14-20 | M | 133 | 32 | 4.10 | 19 | 332 | Intragastric Catheter |
| C15-20 | M | 138 | 46 | 5.40 | 26 | 332 | Intragastric Catheter |
| C20-20 | M | 141 | 32 | 3.91 | 19 | 83 | Intragastric Catheter |
| C48-19 | M | 112 | 45 | 5.00 | 27 | 166 | Intragastric Catheter |
| C49-19 | M | 114 | 29 | 4.40 | 17 | 166 | Intragastric Catheter |

### Acclimation and Surgical Procedures

As in our prior experiments (Patel et al., 2018; Bielanin et al., 2020; Miller et al., 2020), animals were gradually acclimated over at least 5 weeks for 1-hour of confinement in a nylon restraint bag attached to a recording platform using Velcro straps. Subsequently, an aseptic recovery surgery was performed in a dedicated operating suite as described below to attach two-contact platinum-iridium 90/10 electrodes in a silicone patch (Micro-leads Inc., Somerville, MA, USA), similar to those used in prior studies (Nanivadekar et al., 2019; Shulgach et al., 2021), to the proximal duodenum and gastric antrum, on the ventral surface (*Figure 1*). In addition, custom-fabricated electrodes consisting of flexible insulated wire (Cooner Wire, Chatsworth, CA USA) embedded in medical grade silicon (Bentec Medical, Woodland, CA USA) were attached to the diaphragm and abdominal muscles (*Figure 1*).

**Fig. 1.**
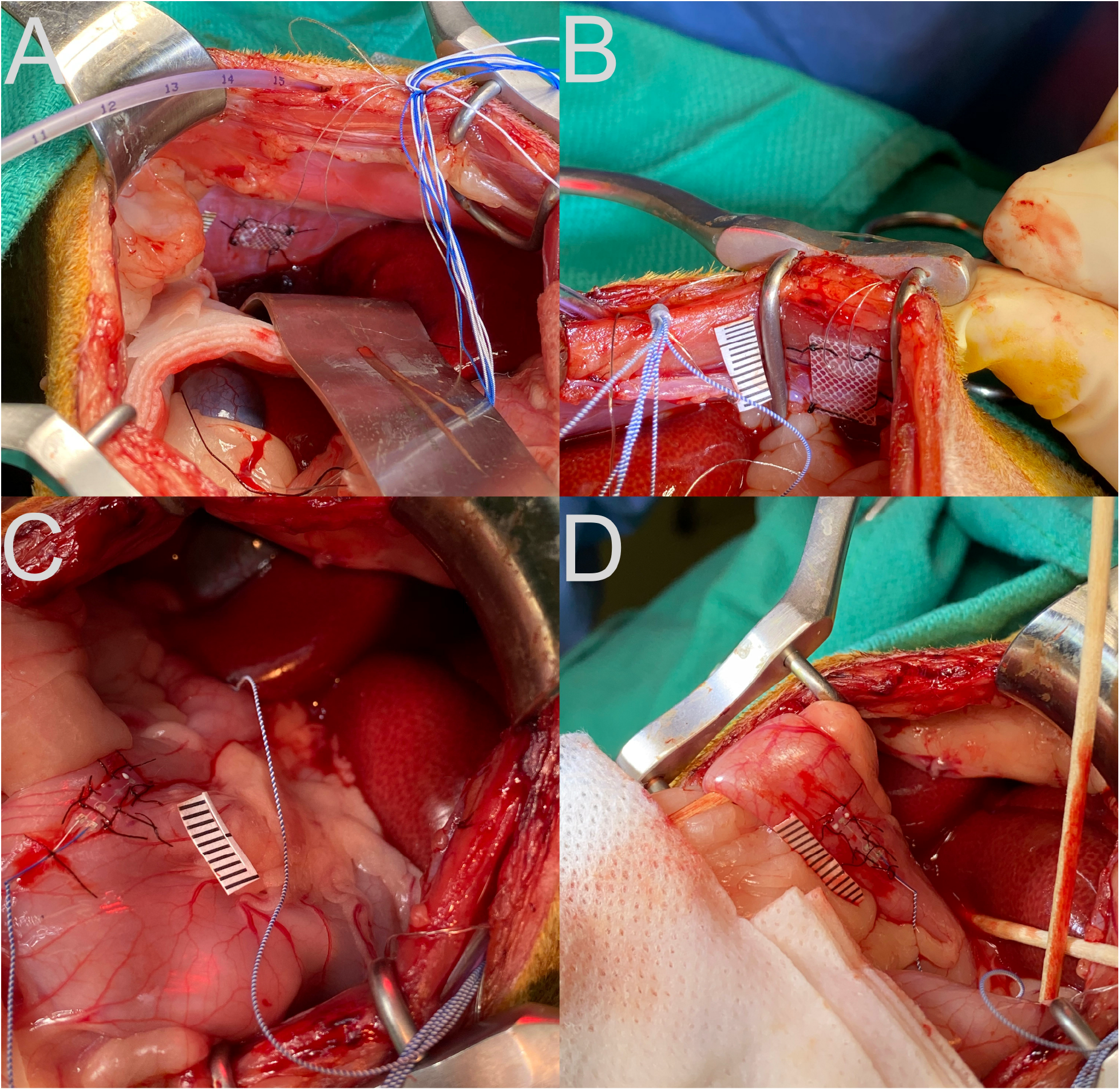
Placement of recording electrodes on the diaphragm (**A**), abdominal musculature (**B**), stomach antrum (**C**), and upper duodenum (**D**). A ruler with 1 mm hatch marks is placed next to the electrode in each panel.

For the surgery, animals were initially anesthetized with an intramuscular injection of ketamine (15 mg/kg) and acepromazine (0.5 mg/kg) to permit the insertion of an endotracheal tube. Subsequently, anesthesia was maintained with 1-2% isoflurane vaporized in oxygen delivered through that tube. Heart rate, respiration rate, and blood oxygen saturation, as well as withdrawal reflexes, were monitored throughout the surgery to guide the concentration of isoflurane that was delivered. A heating lamp and pad were used to keep the animal’s body temperature near 38°C. Saline was administered intravenously throughout the surgery to maintain hydration.

A laparotomy was performed to expose the abdominal organs. Electrodes were secured to the gastric antrum, duodenum, costal diaphragm, and inner surface of the transversus abdominis using sutures through the silicon patch surrounding the recording leads, as shown in *Figure 1*. A small incision was made through the lateral edge of the gastric fundus for introduction of an 8-french intragastric pediatric feeding tube. The tube was secured in place using a purse-string silk suture and Vetbond adhesive (3M, St. Paul, MN USA). The abdominal cavity was lavaged with sterile saline prior to separately closing the abdominal muscles and skin using sutures. Connectors and leads from the electrodes and the distal end of the intragastric catheter were routed subcutaneously to the neck using a trocar. A small area of the skull was exposed, self-tapping screws were inserted, and the connectors were attached to the skull using Palacos bone cement (Heraeus Medical, Hanau, Germany).

A fentanyl transdermal patch (25 μg/h delivery) was attached to the shaved skin on one leg, which provided analgesia for 72 h. In addition, 0.3 mg/kg Meloxicam was provided subcutaneously at the end of surgery and re-administered daily upon veterinary recommendation to alleviate indicators of pain or discomfort. The long-acting antibiotic Convenia (Zoetis, New York, NY USA) was administered during surgery (8 mg/kg subcutaneously).

Following a ^~^2-week post-surgical recovery period, an additional period of acclimation for restraint occurred (see *Table 1*) prior to commencement of data recording.

### Data Recording

During recording sessions, signals from the connectors for the electrodes attached to the diaphragm and abdominal muscles were led into Model 1800 AC differential amplifiers (A-M Systems, Sequim, WA USA), amplified 100 times and filtered with a bandpass of 10-10,000 Hz. Signals from the electrodes positioned on the stomach and intestine were led into Model 3000 AC/DC differential amplifiers (A-M Systems) with the high-pass filter in DC mode, and amplified 50 times. The signals were digitized and recorded at 1 kHz by a Micro1401 mk2 data collection system and Spike-2 version 7 software (Cambridge Electronic Design, Cambridge, UK). Examples of recordings are provided in *Figure 2*.

**Fig. 2.**
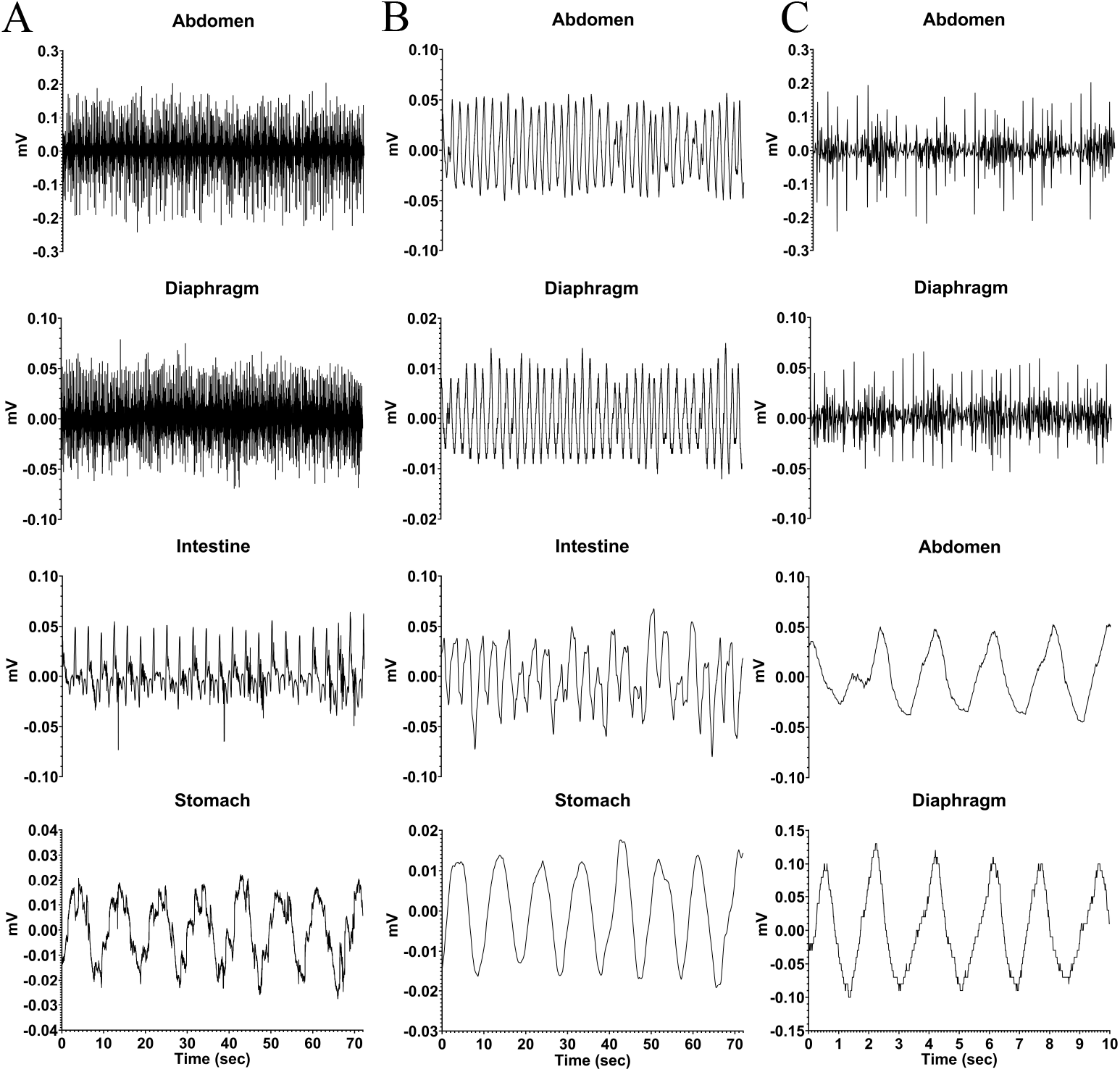
Recordings from one animal (C2-21) during a baseline trial where neither saline nor CuSO_4_ was administered. **A:** raw EMG recordings from the abdominal musculature, diaphragm, intestine (duodenum), and stomach. **B:** same recordings, smoothed with a 0.2 s time constant. **C:** diaphragm and abdominal muscle activity for a shorter time period (first 10 seconds of traces in **A**) so that details of recordings are evident. The top two traces are raw EMG recordings, while the bottom two traces depict the same data smoothed with a 0.2 s time constant.

A number of recording sessions were performed to validate that recordings were stable, and in at least one of those sessions 20 ml of saline was infused into the stomach following an 12-18 hr fast, while recordings from implanted electrodes were made to determine heart rate, respiration rate, and the frequency of stomach and duodenal slow wave activity. Following the collection of these baseline data, a terminal recording session occurred when 83-332 mg of CuSO_4_ dissolved in saline (see *Table 1* for the dose in each animal) was infused into the stomach, through oral gavage in three cases and the intragastric tube in the other seven animals. Animals were fasted overnight prior to this experimental session. Oral gavage occurred in cases where the intragastric catheter became blocked prior to the final experimental session. As during the control session when saline was infused into the stomach, activity recorded from electrodes was used to determine heart rate, respiration rate, and the frequency of stomach and duodenal slow wave activity prior to and following CuSO_4_ administration. Stereotyped contractions of the diaphragm and abdominal musculature were used to determine the timing of emetic episodes (Yates et al., 2014). These data were employed to ascertain whether prodromal changes in physiological parameters predict an emetic episode evoked by CuSO_4_.

During control recording sessions, saline was administered 10 minutes following the onset of the recording, so baseline physiologic parameters could be determined. Recordings continued for 60 minutes. A similar recording strategy was used during the final session when CuSO_4_ was administered. Following the conclusion of recordings in this session, animals were anesthetized by an intramuscular injection of ketamine (20 mg/kg) and acepromazine (0.2 mg/kg) followed by an intraperitoneal injection of pentobarbital sodium (40 mg/kg). Animals were then transcardially perfused with saline followed by 4% paraformaldehyde fixative. The placement of recording electrodes was evaluated by dissecting the abdominal cavity and extracting the diaphragm and GI tract for imaging.

### Data Analysis

Offline analysis of recordings was performed using Spike-2 version 9, MATLAB (Mathworks, Natick, MA USA), and Python (Python Software Foundation) software. As an initial step, DC drift was removed from all traces. Signals from respiratory muscles were smoothed (time constant of 0.2 s) and/or rectified for some analyses. Myoelectric signals recorded from the stomach and intestine electrodes were processed using a bandpass 2nd order Butterworth filter with a 0.05-0.5 Hz (3-30 cycles per minute, cpm) cutoff range, and then downsampled to 10Hz. Because the head was not restrained, some non-physiological high amplitude artifacts lasting 1-20 seconds were observed during head movements. These artifacts were cleaned by implementing a blanking window around the artifact period with 1 second before and after the event and replacing the blanked portion with a linearly interpolated line that spanned the window. The power spectrum for myoelectric signals was calculated by computing the fast Fourier transform (FFT, 0.014 cpm bin size). The dominant frequency was determined by finding the median frequency of a power spectral density estimate at each one-minute segment throughout the length of the signal. Statistical analyses were conducted and some figures were plotted using Prism 9 software (GraphPad software, San Diego, CA USA). Data are presented as means ± one standard deviation.

## 3 Results

*Figure 2* shows raw and processed data recorded during a baseline run in one animal. *Column A* illustrates raw traces, and in *Column B* the same traces were smoothed (time constant of 0.2 s). *Column C* expands the first 10 seconds of respiratory muscle activity shown in *column A* so that details of traces are evident; the top two traces show raw activity, and the bottom two are smoothed activity (time constant of 0.2 s). Both the electrocardiogram and respiratory muscle electromyographic (EMG) activity were evident in recordings from the diaphragm and abdominal muscles, such that these traces could be used to determine both heart rate and respiration rate. In addition, slow wave activity was evident for both the stomach (frequency of ^~^0.1 Hz or 6 cpm) and duodenum (frequency of ^~^0.3 Hz or 18 cpm), which occurred at a different rate than respiration (frequency of ^~^0.6 Hz or 36 cpm) indicating it was not an artifact related to abdominal movements during breathing. Heart rate, respiration rate, and frequency of stomach and duodenal slow wave activity are shown in *Table 2*, determined for each animal during a 60-minute period of a baseline recording. Some physiologic parameters could not be determined in a few animals due to low-quality recordings from particular electrodes. The average baseline heart rate for all animals was 163 ± 26 beats per minute, and the average respiratory rate was 34 ± 16 breaths per minute. Slow wave activity occurred at 6 ± 1 cpm in the stomach and 20 ± 1 cpm in the duodenum.

**Table 2.**

| **Animal ID** | **Average Heart Rate (BPM)** | **Average Respiration Rate (BPM)** | **Average Intestine Slow Wave Activity (cpm)** | **Average Stomach Slow Wave Activity (cpm)** |
| --- | --- | --- | --- | --- |
| C1-21 | 160 | x | x | x |
| C2-21 | 163 | 32 | 19 | 5 |
| C3-21 | 232 | 73 | 20 | 8 |
| C11-21 | 150 | 23 | 18 | 7 |
| C13-20 | 139 | 41 | 19 | 4 |
| C14-20 | 161 | 21 | 18 | 6 |
| C15-20 | 145 | 22 | 19 | 6 |
| C20-20 | 142 | 20 | 23 | 6 |
| C48-19 | 182 | 47 | 21 | 6 |
| C49-19 | 153 | 30 | 20 | 5 |

Baseline physiological measurements for each animal.

Following the administration of CuSO_4_, we visually observed the animals for indications of retching and compared these observations to recordings from the electrodes. In all cases, a retching period occurred as a series of contractions that was accompanied by large voltage signals recorded from both the diaphragm and abdominal muscles, as shown in *Figure 3*. The left panels illustrate raw respiratory muscle EMG activity, whereas the traces are rectified and smoothed (time constant of 0.2 s) in the right panels. Respiratory-related activity of the muscles is evident prior to and following the retching period. These stereotyped responses served as markers for the onset and offset of an emetic episode.

**Fig. 3.**
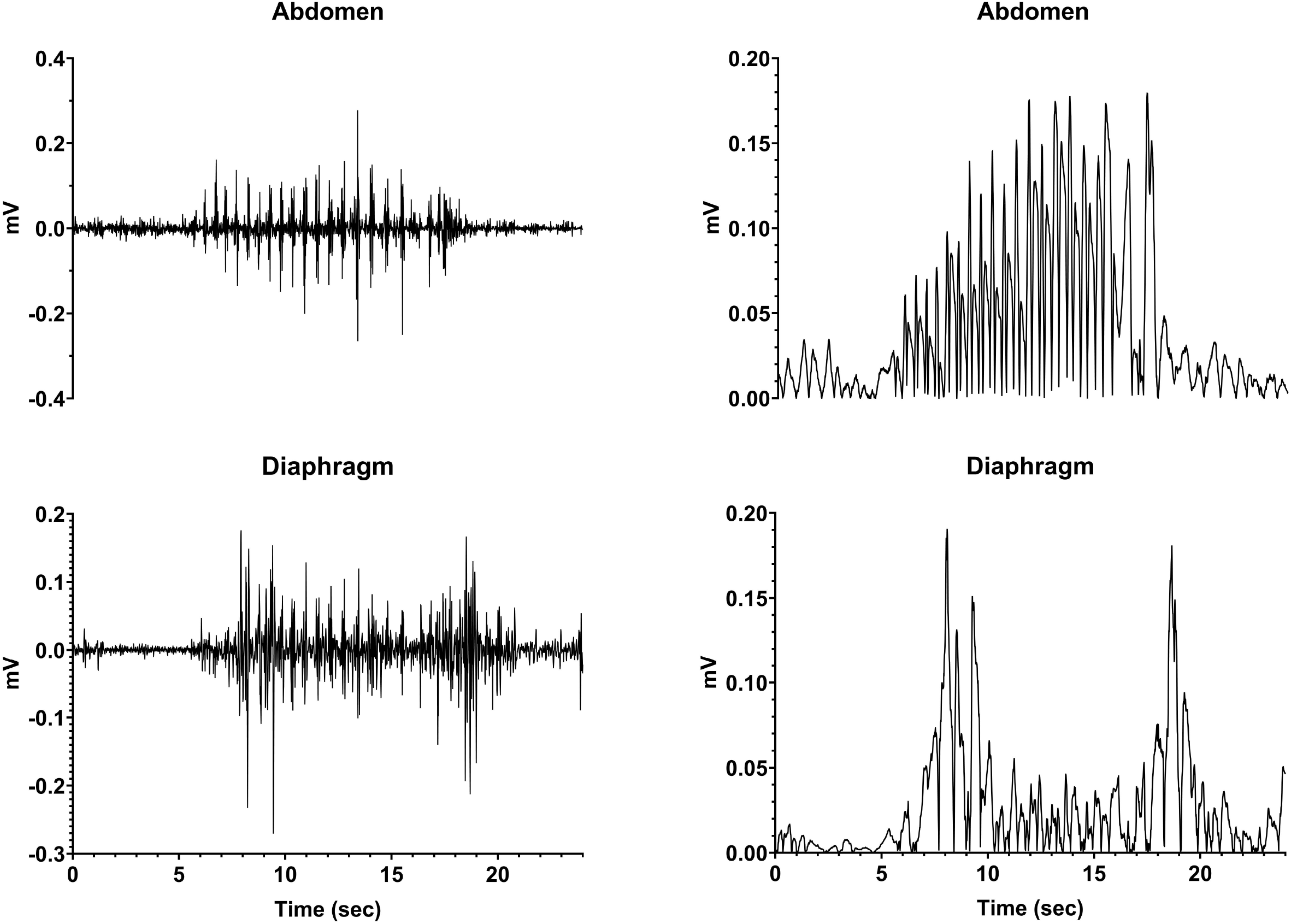
Recordings of diaphragm and abdominal muscle activity during a retching episode (continuous series of retches). The left traces are raw EMG activity, whereas the recordings are rectified and smoothed (time constant of 0.2 s) in the right panels.

*Table 3* indicates the responses of the animals to CuSO_4_ administration. All animals exhibited a retching period, and in six of ten cases infusion of CuSO_4_ produced a retching response within 10 minutes. A linear regression analysis showed no evidence that the CuSO_4_ dose affected the time to the first retch (P=0.58; R^2^=0.04), the duration of the first retching episode (P=0.48; R^2^=0.06), or the number of emetic responses that were elicited (P=0.32; R^2^=0.12). Although numbers are small, there were also no overt differences in the emetic responses of animals of different sexes to CuSO_4_ administration.

**Table 3.** Responses of animals to CuSO_4_ administration.

| **Animal ID** | **Sex** | **Route of CuSO<sub>4</sub> Administration** | **CuSO<sub>4</sub> Dosage (mg)** | **Time to First Retch (min)** | **Duration of First Retching Episode (sec)** | **# of Emetic Episodes Elicited** |
| --- | --- | --- | --- | --- | --- | --- |
| C1-21 | F | Intragastric | 166 | 25 | 11.2 | 3 |
| C2-21 | F | Oral | 332 | 4 | 7.2 | 4 |
| C3-21 | F | Oral | 332 | 33 | 5.3 | 2 |
| C11-21 | F | Oral | 332 | 6 | 8.9 | 2 |
| C13-20 | M | Intragastric | 83 | 14 | 11.8 | 2 |
| C14-20 | M | Intragastric | 332 | 3 | 7.5 | 8 |
| C15-20 | M | Intragastric | 332 | 33 | 13.5 | 3 |
| C20-20 | M | Intragastric | 83 | 6 | 10.1 | 2 |
| C48-19 | M | Intragastric | 166 | 2 | 7.0 | 4 |
| C49-19 | M | Intragastric | 166 | 10 | 6.2 | 3 |

Changes in heart rate, respiration rate, and the frequency of slow wave activity in the duodenum and stomach elicited by saline and CuSO_4_ administration in one animal (C2-21) are shown in *Figure 4*. Each data point indicates averaged data collected over a 1-minute interval. In this animal and all others, saline administration did not elicit retching. However, within a few minutes of CuSO_4_ administration a retching period occurred, as denoted by a gold area and black vertical lines. Just prior to retching, a distinct increase in heart rate and frequency of intestinal slow-wave activity occurred. Heart rate then dropped at the onset of the retching episode, whereas the increased frequency of intestinal slow-wave activity was sustained throughout the retching period and a few subsequent minutes of recording. In addition, respiratory muscle activity decreased prior to retching and the frequency of stomach slow wave activity increased, although the baselines were less stable than for heart rate and intestinal slow wave activity, so these responses are less evident.

**Fig. 4.**
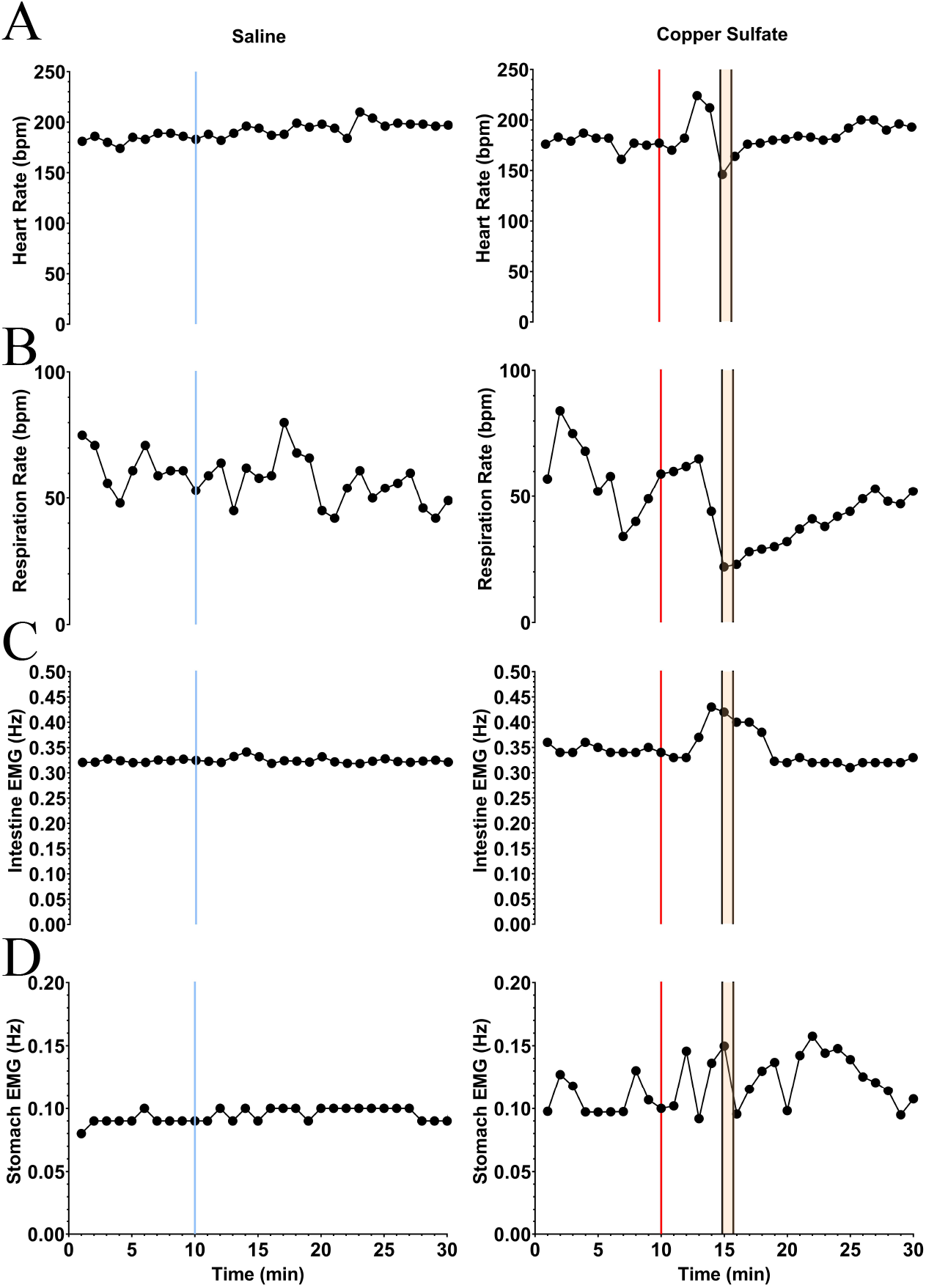
Changes in heart rate, respiration rate, and the frequency of slow wave activity in the duodenum and stomach elicited by saline (*left column*) and CuSO_4_ (*right column*) administration in one animal. Each data point indicates averaged data over a 1-s period. The vertical blue line in the left column denotes the time of saline administration, whereas the vertical red line in the right column shows when CuSO_4_ was provided. The time period when retching occurred following CuSO_4_ administration is designated by gold shading and black vertical lines.

To enhance visualization of changes in the frequency of stomach and duodenal muscle slow-wave activity, one-minute averages of data were also displayed as spectrograms generated using MATLAB, as illustrated in *Figure 5*. Data for the same animal are provided in *Figures 4 and 5*, with a longer time period depicted in *Figure 5*. These spectrograms indicate that little change in slow wave activity occurred following intragastric saline administration, but a prominent change in duodenal slow wave activity commenced before and extended through the first retching period (indicted by vertical black lines). Alterations in stomach slow wave activity are also evident but are less prominent due to variability in the baseline. A second period of retching occurred approximately 30 minutes following the first episode (designated by a second group of black bars) but was not accompanied by appreciable changes in the frequency of slow wave activity recorded from the stomach or duodenum.

**Fig. 5.**
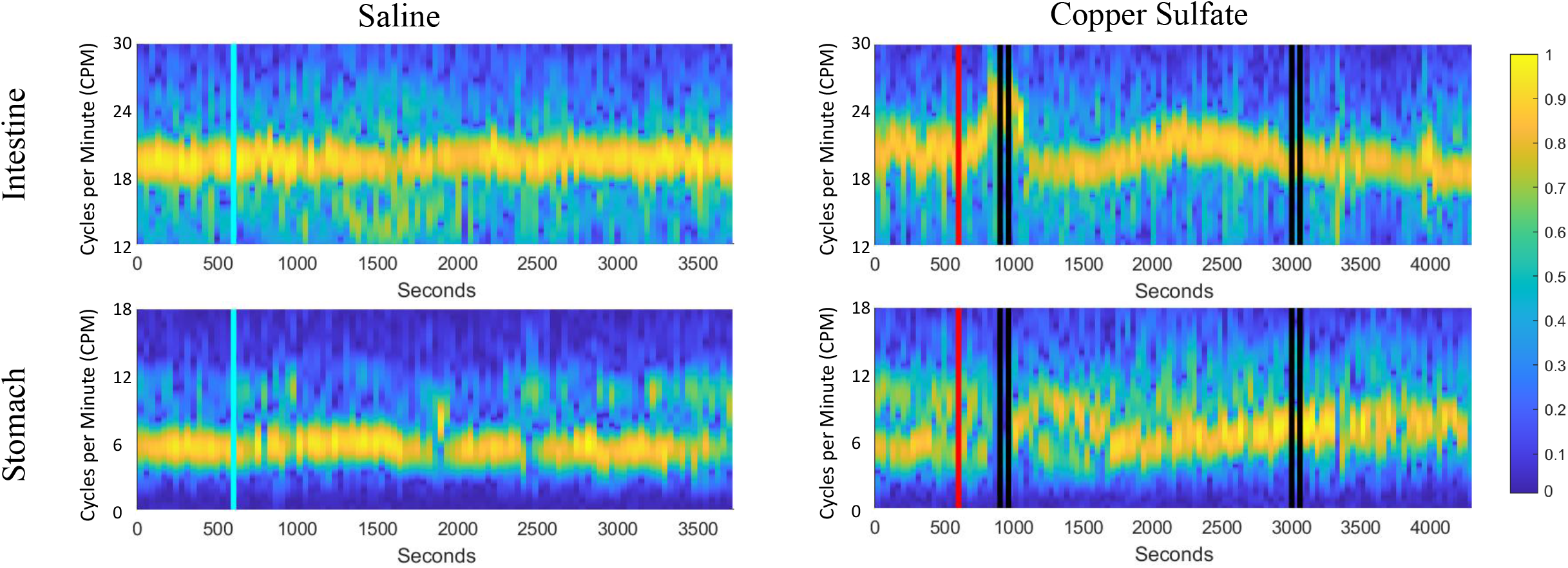
Spectrograms of duodenal and stomach slow-wave activity following administration of saline (*left column*) and CuSO_4_ (*right column*). Yellow represents the dominant frequency and blue the least dominant (scale of normalized signal power, 0 to 1). As in Fig. 4, the blue vertical line indicates saline administration, the red vertical line designates CuSO4 administration, and the black vertical lines show when retching occurred.

*Figure 6* compares average heart rate, respiration rate, and the span of normal intestinal and stomach slow-wave activity (normogastric activity) during the baseline period (10-minute period prior to CuSO_4_ administration), and the 10-minute period during which the first retching episode occurred. As in our previous study (Nanivadekar et al., 2019), normogastric activity was defined by establishing a 99% confidence interval of EMG activity during the baseline period. Average EMG activity during each minute of both the baseline and retching periods was classified as normogastric (within the confidence interval), tachygastric (above the confidence interval), or bradygastric (below the confidence interval). As indicated in *Table 3*, six of the animals retched within 10 minutes of CuSO_4_ administration. For these animals, whose data are indicated in green in *Figure 6*, the retching period was defined as the 10-minute epoch immediately following CuSO_4_ delivery. In the four animals with delayed retching responses, whose data are indicated in red in *Figure 6*, the retching period was defined as the 10-minute epoch preceding the final retch of the initial emetic response. Due to deterioration of recordings from some electrodes over time, data are missing from some animals. Heart rate could be determined for every animal, whereas respiration rate and intestinal EMG activity was evident in seven animals, and stomach EMG activity in six animals. Two-tailed paired t-tests were used to compare physiologic parameters in the baseline and retching periods; p values are provided above each panel.

**Fig. 6.**
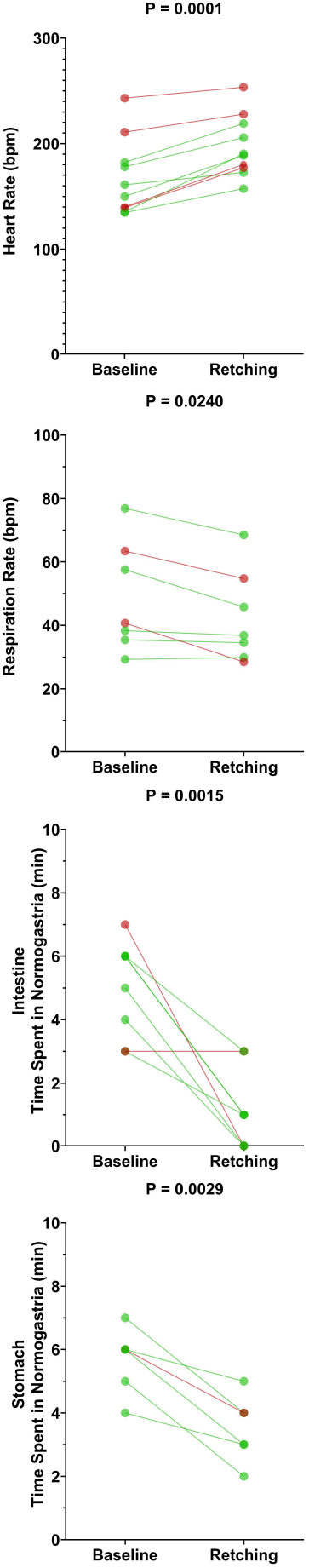
Effects of CuSO_4_ administration on physiologic parameters associated with the first retching episode. Each data point represents an average of data over 10 minutes during the baseline (prior to CuSO_4_ administration) or retching periods. For the six animals that retched within 10 minutes of providing CuSO_4_ (data indicated by green lines and symbols), the retching period was defined as the 10 minutes following CuSO_4_ administration. For the four animals where retching occurred at longer latency (data indicated by red lines and symbols), the retching period was defined as the 10 minutes prior to the final retch of the initial episode. P values above each panel are for comparisons between each physiologic parameter during the baseline and retching periods (two-tailed paired t-test).

CuSO_4_ administration produced a significant increase in heart rate (p=0.0001) and a decrease in respiratory activity (p=0.024). Heart rate increased by an average of 19 ± 11%, and respiratory activity decreased by an average of 11 ± 11%. Both intestinal and stomach EMG activity were significantly disrupted (less time spent in normogastria) by CuSO_4_: intestinal normogastric activity decreased by 73 ± 34% (p=0.0015), whereas stomach normogastric activity decreased by 38 ± 16% (p=0.0029). The disruption in intestinal EMG activity was particularly striking in the five animals where baseline activity was stable (normogastric ≥ 40% of the baseline period). In these cases, intestinal normogastric activity decreased by 86 ± 19% following CuSO_4_ administration.

A similar analysis was conducted for intragastric saline administration to the animals. Since saline did not evoke retching in any of the animals, the effects of saline on physiologic parameters were determined during the epoch defined as the retching period following CuSO_4_ administration. A two-tailed paired t-test showed no effects of saline administration on heart rate (p=0.63), respiration rate (p=0.83), or disruption of normogastric intestinal (p=0.61) or stomach (p=0.60) activity.

## 4 Discussion

This study incorporated extensive monitoring of physiologic parameters in an alert animal model during the administration of an agent that produces rapid emetic events, allowing the analysis of prodromal physiologic changes that precede vomiting. The frequency of baseline GI myoelectric activity noted in this study is similar to that documented in prior experiments (Connor, 1979; Roche et al., 1982; Lang, 1999). CuSO_4_ infusion elicited a significant increase in heart rate, decrease in respiratory rate, and a disruption of gastric and intestinal EMG activity several minutes prior to emesis. The most striking and consistent prodromal physiologic changes were an increase in heart rate and a reduction of time that duodenal activity was normogastric. In contrast to the prevailing focus of work showing changes in the gastric EMG signals after emetic stimulation (Koch et al., 1990; Uijtdehaage et al., 1992; Kiernan et al., 1997; Parkman et al., 2003), our work shows that EMG activity changes were most consistent in the duodenum. It is postulated that intestinal EMG activity was related to a retrograde movement of chyme from the intestine to the stomach, which could be associated with the prodromal retrograde contractions reported after emetic treatments (Lang, 1990 #29004; Ueno and Chen, 2004; Koch, 2014; Heyer et al., 2018).

Administration of saline did not induce similar physiologic changes as CuSO_4_, indicating that they were not induced by a distension of the stomach with fluid. Increasing the dose of CuSO_4_ did not alter the physiologic effects of the treatment or the length of time between the administration of the agent and the first retching event. Systematic changes in heart rate and disruption of GI myoelectric activity only occurred within a short period preceding and during emesis (see *Figure 4*). These observations suggest that the changes in heart rate and GI myoelectric activity observed in the study were related to nausea, and not nonspecific actions of CuSO_4_ such as irritation of the stomach lining.

Several caveats and limitations of the current study should be noted. First, in some cases we were not able to administer CuSO_4_ through the implanted gastric tube, and instead gavaged CuSO_4_ solution. Although the physiologic effects in these animals were similar to those in animals that received the agent intragastrically (see *Table 3*), it is possible that the distribution of CuSO_4_ in the stomach differed between the routes of administration. Moreover, in general there was variability of stomach geometry and placement of the tip of the delivery tube in the gastric compartment; therefore, it is difficult to determine the specific chemical placement and timing of movement of the emetic agent along the GI tract. In addition, we also did not attempt to replicate the CuSO_4_ tests within each animal because CuSO_4_ is known to damage the GI mucosal surface (Chuttani et al., 1965; James et al., 1999; Nastoulis et al., 2017).

This issue of testing repeatability is likely important to translate these findings for potential clinical application. Although acute emesis (and nausea) can be controlled by administration of currently available antiemetics, such as 5-HT3 and NK-1 receptor antagonists, the chronic nausea and vomiting that occurs in GI disease is more difficult to control (Hasler et al., 2017; Lacy et al., 2018). Potentially, these chronic clinical conditions could be modeled with a repeated testing paradigm using pre-clinical models. Importantly, this work also needs to be extended to include other emetic stimuli, acting on non-vagal pathways, including motion and those that activate area postrema. Nonetheless, the current findings provide a blueprint for these future studies by highlighting the importance of monitoring intestinal EMG activity. It is also possible that a machine learning paradigm that incorporates changes in heart rate and disruption of intestinal myoelectric activity would provide a particularly effective method of predicting the onset of emesis.

## Acknowledgments

The authors thank Brandon King, Henry Ramos Acosta, and Robert Khorami for their assistance with data collection and analysis. A preliminary report of these findings has been published on the bioRxiv preprint server and as a poster presentation at the 2022 Experimental Biology meeting in Philadelphia, PA.

